# Improved Tumor Blood Flow Enhances the Abscopal Effect: Preclinical Assessment in Mice Treated with Combined Radiation and PD-1 Blockade Therapy

**DOI:** 10.1101/2025.05.04.652150

**Authors:** Kota Yamashita, Jeffrey R. Brender, Yu Saida, Yasunori Otowa, Horie Kazumasa, Takeshi Ito, Kazutoshi Yamamoto, Helmut Merkle, W. M. Linehan, Murali C. Krishna, Shun Kishimoto

**Affiliations:** Radiation Biology Branch, Center for Cancer Research, National Cancer Institute, NIH, Bethesda, Maryland, United States; Clinical Cancer Metabolism Facility, National Cancer Institute, NIH, Bethesda, Maryland, United States; Department of Respiratory Medicine and Infectious Diseases, Niigata University Graduate School of Medical and Dental Sciences, Niigata, Japan; Urologic Oncology Branch, Center for Cancer Research, National Cancer Institute, NIH, Bethesda, Maryland, United States; Laboratory for Functional and Molecular Imaging, National Institute of Neurological Disorders and Stroke, National Institutes of Health, Bethesda, MD, USA

**Keywords:** abscopal effect, PD-1 blockade, radiation therapy, carbogen

## Abstract

The abscopal effect, where localized radiation therapy induces regression of distant metastatic lesions through immune activation, shows promise for treating metastatic cancer but occurs inconsistently. Here we demonstrate that tumor perfusion critically influences systemic immune responses to combination therapy with radiation and PD-1 blockade. Using multimodal imaging including DCE-MRI, EPR oximetry, and hyperpolarized 13C-MRI, we show that successful abscopal responses in MC38 tumors are characterized by enhanced perfusion, reduced hypoxia, decreased cellularity, and lower glycolytic activity in remote tumors. Notably, pre-treatment perfusion metrics (AUC1min) and extracellular volume (AUC10min) in primary tumors predict subsequent growth of remote tumors, while the same measurements in remote tumors lack predictive value. Based on these findings, we enhanced the abscopal effect by exposing mice to carbogen (95% O2 + 5% CO2) during radiation therapy. Carbogen exposure increased tumor perfusion by 71% (AUC1min) and significantly improved systemic responses in the checkpoint blockade responsive MC38 model but not in the poorly responsive B16.F10 tumors. The enhanced response correlated with increased activation of CD8^+^ T cells in tumor-draining lymph nodes and elevated serum HMGB-1 levels. RNA sequencing revealed significant extracellular matrix remodeling in carbogen-treated tumors. These results establish tumor perfusion as both a predictive biomarker and a modifiable determinant of systemic immune responses, suggesting that perfusion-based patient stratification and vascular modification strategies could improve outcomes in combination immunotherapy and radiation treatment.

## Introduction

The abscopal effect is a rare but notable phenomenon where radiation therapy (RT) induces regression in metastatic lesions distant from the irradiated site.(1,2) Its occurrence with RT alone is so rare that it has historically been a subject of debate within the scientific community.(3) This rarity is largely attributed to the immunosuppressive characteristics of the tumor microenvironment (TME), which hinder effective systemic immune activation.(4) In recent years combining RT with immunotherapy, particularly immune checkpoint inhibitors, has led to an increase in documented cases of the abscopal effect.(4–6) However, even with these advancements, the effect remains inconsistent, and its full mechanism has yet to be elucidated.

This inconsistency reflects the fact that the abscopal effect is not universally observed but instead depends on specific situational factors related to both the treatment approach and the patient’s tumor biology. Radiation therapy can trigger an immune response by exposing tumor antigens, effectively turning the irradiated lesion into a “vaccine” that primes the immune system to attack distant metastases.(5) However, this process is frequently hindered by the immunosuppressive TME, which creates significant barriers to effective immune activation.(7) The addition of immunotherapy, such as immune checkpoint inhibitors, has been shown to mitigate some of these barriers by enhancing T cell activity and reducing immune suppression. Nevertheless, variability in patient-specific factors and tumor characteristics continues to make the effect unpredictable.(8)

Moreover, the high radiation doses often required to induce an abscopal effect carry risks of adverse events, including damage to healthy tissues and long-term complications.(5,9) These risks underscore the urgent need for reliable biomarkers that can predict which patients are most likely to benefit from RT-induced systemic responses.(5,10) Such biomarkers could enable clinicians to tailor treatment strategies more effectively, minimizing unnecessary exposure to high-dose radiation while maximizing therapeutic benefits.

A number of biomarkers have been proposed to assess immune responses in cancer therapy,(10) including p53 mutations,(11) total lymphocyte counts,(12,13) markers for DNA repair,(14) tumor neoantigens,(2,15) tumor infiltrating lymphocytes,(16–18) myeloid-derived suppressor cells (MDSCs),(19) and signaling molecules like damage-associated molecular patterns (DAMPs) and cytokines.(20) Despite their potential utility, each of these biomarkers faces significant methodological and practical limitations that restrict their reliability and clinical applicability. For instance, assessing infiltrating lymphocytes in the TME requires invasive tissue sampling followed by labor-intensive spatial analysis. This process is not only time-consuming but also prone to interobserver variability. Similarly, measuring signaling molecules such as DAMPs and cytokines presents challenges. Cytokine levels are difficult to quantify accurately because of their short half-life and low baseline concentrations in serum, while DAMPs are released in response to various types of cellular stress and injury, not just those associated with cancer. Both often lack specificity since they are modulated by a wide range of conditions beyond cancer therapy. In addition to these practical challenges, many biomarkers, including lymphocyte counts, suffer from inherent biological variability. These counts fluctuate naturally and can be influenced by unrelated factors such as infections or medications. These issues underscore the need for biomarkers that are not only precise and reproducible but also capable of capturing the complex dynamics of immune responses within the TME.

Among these situational factors influencing treatment outcomes, metabolic dynamics within the TME play a particularly critical role. Tumor cells often dominate glucose metabolism through enhanced glycolysis, depriving cytotoxic CD8^+^ T cells of the energy required for optimal function. This metabolic imbalance weakens T cell-mediated anti-tumor responses and contributes to immune suppression. Hypoxia within the TME further exacerbates this suppression by impairing mitochondrial function in T cells, leading to their exhaustion and diminished efficacy against distant metastases.(21,22) Immune checkpoint inhibitors like anti-PD-1 can partially counteract these effects by reducing cancer cell glycolysis and reactivating T cells.(23,24) However, variability in hypoxia levels and vascular perfusion across tumors underscores why treatment outcomes remain inconsistent and highly situational.

Imaging techniques such as dynamic contrast-enhanced (DCE) MRI, Electron Paramagnetic Resonance Imaging (EPRI), and hyperpolarized ^13^C MRI have emerged as valuable tools for assessing these characteristics.(25–28) These modalities allow for detailed evaluations of perfusion and oxygenation within tumors - factors closely linked to immune cell infiltration and activation. For example, DCE MRI can quantify vascular perfusion and permeability, while EPRI provides precise measurements of tumor oxygenation levels. Together, these imaging approaches offer a comprehensive view of how metabolic and functional changes within tumors may influence therapeutic outcomes.

Our study utilized these imaging techniques to investigate tumor models treated with a combination of RT and PD-1 blockade. We found that higher baseline blood perfusion in primary tumors was strongly associated with a more robust abscopal response. Importantly, we demonstrated that carbogen exposure during RT enhanced tumor perfusion without requiring additional radiation doses, thereby improving the likelihood of inducing an abscopal effect while mitigating potential adverse effects of high-intensity RT. This approach not only increased oxygen delivery but also facilitated the release and uptake of tumor antigens necessary for effective immune activation.

## Materials and Methods Mice and cell lines

All animal experiments were conducted according to a protocol approved by the Animal Research Advisory Committee of the NIH (RBB-159-3E) in accordance with the National Institutes of Health Guidelines for Animal Research. Female C57BL/6 (B6) mice were obtained from the Frederick Cancer Research Center, Animal Production, and housed in a specific pathogen-free environment under controlled conditions (22°C–24°C, 40%– 60% humidity, 12-hour light/dark cycle). They were used at 8–12 weeks of age. The study utilized MC38 colon adenocarcinoma and B16-F10 melanoma cell lines to represent immunogenic and less immunogenic models, respectively, based on their relevance to preclinical studies of immune checkpoint blockade therapy. MC38 cells were purchased from Kerafast, while B16-F10 cells were sourced from our lab’s frozen stock, tested in February 2020, and authenticated by IDEXX RADIL using microsatellite markers. Molecular testing for multiple pathogens, including Mycoplasma, was performed upon receipt and before in vivo studies. Both cell lines were cultured in DMEM with 4.5 g/L glucose, 10% FCS, and 1% Hyclone penicillin/streptomycin solution (10,000 units each) from Cytiva.

### Tumor models

To establish a tumor model demonstrating the abscopal effect induced by RT and PD-1 inhibitor combination therapy, MC38 or B16-F10 tumor cells were inoculated subcutaneously into both legs of C57BL/6 mice under sterile conditions. For the MC38 model, 1x10^6^ and 2x10^5^ MC38 tumor cells were inoculated subcutaneously into right (primary tumor) and left (remote tumor) hindlegs of C57BL/6 mice to form a smaller remote tumor, respectively. For the B16.F10 model, 2.5 x 10^5^ and 0.5 x 10^5^ B16.F10 tumor cells were inoculated into right and left hindlegs of C57BL/6 mice, respectively.

### In Vivo PD-1 Antibody and Radiotherapy Treatment

Four treatment groups were prepared: IgG isotype antibody control, irradiation alone, αPD-1 antibody treatment alone, and combination therapy with RT and αPD-1 antibody treatment. Tumor-bearing mice received intraperitoneal injections of 200 μg αPD-1 antibody or isotype control antibodies (BioXcell) on day 0, 3, and 6 post-inoculation when tumors reached approximately 100 mm³ as measured by calipers (length × width²/2). This was followed by three consecutive daily irradiations of 8 Gy delivered locally to the primary tumor using an X-RAD 320 irradiator with lead shielding to protect surrounding tissues. Imaging experiments were conducted pre-treatment on the irradiated primary tumor in the right hind leg and on the unirradiated remote tumor in the left hind leg nine days after first antibody injection.

### Flow cytometry

To examine immune activation in tumors and tumor-draining lymph nodes (TDLNs), treated mice were euthanized by cervical dislocation, and tissues were harvested to prepare single-cell suspensions. TDLNs were mechanically dissociated. Solid tumors were enzymatically dissociated with a mixture of 0.1% collagenase, 0.01% DNase I, and 2.5 U/mL hyaluronidase for 3 hours at 37°C with gentle agitation. After digestion, the cell suspensions were filtered through a 70 μm cell strainer to remove debris and washed with HBSS.

For the analysis of tumor-infiltrating lymphocytes (TILs), single-cell suspensions from tumors were stained with FITC-conjugated anti-mouse CD4, APC-conjugated anti-mouse CD3, PE-conjugated anti-CD8 and Zombie Aqua Fixable Viability Kit (all from BioLegend). Cell surface phenotypes were determined by direct immunofluorescence staining, and data acquisition was performed using a Cytek Aurora (Cytek). Data analysis was conducted using FlowJo software. TILs were identified by gating on live cells based on forward scatter versus side scatter plots.

To assess immune activation in TDLNs, single-cell suspensions were obtained from the right inguinal and iliac lymph nodes. Analysis was performed similarly but cells were stained with PE/Cyanine7 anti-mouse CD69 antibodies (BioLegend) and BUV395 anti-mouse CD25 antibody (BD Biosciences) in addition to the antibodies used in TILs.

### EPR Oximetry

Technical details of the Electron Paramagnetic Resonance Imaging (EPRI) scanner and oxygen image reconstruction were previously described.{Matsumoto, 2006 #68}. Parallel coil resonators tuned to 300 MHz were used for EPRI. After an animal was placed in the resonator, an oxygen-sensitive paramagnetic trityl radical probe, OX063 (1.125 mmol/kg bolus), was injected intravenously under isoflurane anesthesia to ensure consistent physiological conditions and reduce motion artifacts.

Time-domain EPR signals were recorded at ambient temperatures using a single excitation pulse sequence with a pulse width of 60 ns (flip angle ∼50°), repetition time (TR) of 8 μs, and free induction decay (FID) signal averages of 4000. Each signal acquisition included four phase cycles, and 1000 sampled points were recorded with a dwell time of 5 ns. The 3D EPR imaging data were acquired in single-point imaging (SPI) mode, a phase-encoding approach, using a 19 × 19 × 19 Cartesian grid. Raw k-space data underwent correction for DC shifts, and high-frequency noise was filtered using a Tukey window (r = 0.7). Image reconstruction was conducted through fast Fourier transformation after zero-filling the k-space matrix to 64 × 64 × 64 or higher.

The FID signals, lasting between 1–5 ms, were used to generate a series of T2* maps (EPR line width maps), which linearly correlate with the local concentration of oxygen when the concentration of OX063 is low enough to avoid self-line broadening effects. This allowed pixel-wise estimation of tissue partial pressure of oxygen (pO_2_). The repetition time of the measurements was optimized to minimize signal loss while maintaining sufficient sensitivity for oxygen quantification.

After EPRI measurements, corresponding anatomical T2-weighted MR images were collected on a 1 T scanner (Bruker BioSpin MRI GmbH) to provide structural context for the functional oxygen maps. The MR images were co-registered with EPRI data for accurate localization of oxygenation changes within tumors.

### DCE-MRI

Dynamic contrast-enhanced (DCE) MRI studies were performed on a 3 T scanner (Bruker BioSpin MRI GmbH) to assess tumor perfusion and permeability. T1-weighted fast low-angle shot (FLASH) images were acquired with the following parameters: repetition time (TR) = 156 ms, echo time (TE) = 4 ms, flip angle = 45°, four slices, spatial resolution = 0.44 × 0.44 mm, acquisition time per image = 15 seconds, and 45 repetitions. A Gd-DTPA solution (4 mL/g body weight of 50 mmol/L Gd-DTPA) was injected through a tail vein cannula 1 minute after the start of the dynamic FLASH sequence to ensure consistent timing of contrast administration.

To determine local Gd-DTPA concentrations, T1 maps were calculated from three sets of Rapid Imaging with Refocused Echoes (RARE) images acquired before the FLASH sequence. The RARE images were obtained with TR values of 300, 600, 1,000, and 2,000 ms. These T1 maps were used to quantify changes in signal intensity over time, allowing for the calculation of perfusion-related metrics such as the area under the curve at 10 minutes (AUC10min). AUC10min was used as a surrogate marker for extracellular volume at equilibrium.

Motion artifacts during imaging were minimized by anesthetizing mice with isoflurane and securing them in a custom-built holder to maintain consistent positioning throughout the scan.

### ADC Mapping by Diffusion MRI

Diffusion-weighted MRI (DW-MRI) was performed to measure apparent diffusion coefficient (ADC) values as a marker of tumor cellularity. DW-MRI scans were acquired using the same geometry as DCE-MRI to ensure spatial correspondence between datasets. The sequence parameters included five b-values: 12, 50, 500, 800, and 1,500 s/mm²; TR = 1,500 ms; TE = 35 ms; and six averages to reduce motion artifacts. Mice were anesthetized with isoflurane during imaging to minimize motion-related variability.

ADC maps were generated using a custom MATLAB program that fit b-values to the Stejskal-Tanner equation. The ADC values reflect the microscopic motion of water molecules within tissues and are inversely correlated with tumor cellularity. Higher ADC values indicate reduced cellularity due to increased extracellular space or cell death, while lower ADC values suggest higher cellular density.

To ensure data quality, all scans underwent visual inspection for motion artifacts prior to analysis. Regions of interest (ROIs) were manually delineated on tumor areas in each slice using anatomical reference images from DCE-MRI.

### Hyperpolarized 13C MRI

Hyperpolarized [1-13C] pyruvic acid was prepared using a SpinAligner DNP polarizer. Briefly, 18 μL of [1-13C] pyruvic acid containing 25 mmol/L Ox063 was polarized until reaching 80% of the plateau value. The hyperpolarized sample was then rapidly dissolved in 3.2 mL of a superheated alkaline buffer consisting of 40 mmol/L HEPES, NaOH, and 100 mg/L EDTA. The resulting hyperpolarized [1-^13^C] pyruvate solution was intravenously injected into mice through a tail vein catheter at a dose of 12 mL/g body weight to ensure consistent delivery.

Hyperpolarized ^13^C MRI studies were performed on a 3 T Bruker Biospec animal scanner using a home-built ^13^C-1H dual tuned coil. The transceiver coil consisted of two conventional 47 mm long crossed saddle coils with an inner diameter of 31 mm which fits around the mouse bed of the BRUKER 3T system. The conductors were made from silverplated and insulated multistranded AWG 14 or AWG 16 copper wires (homemade Litz wires) to minimize inductance and to maximize Q. Both the ^1^H and ^13^C frequencies were tuned and matched, and anatomical images were acquired after shimming on the proton signal to ensure optimal field homogeneity. Chemical Shift Imaging (CSI) was performed every 12 seconds for a total duration of 240 seconds, focusing on the center of the leg tumors. The imaging parameters included a repetition time (TR) of 1,000 ms, spectral width of 3,300 Hz, flip angle of 10°, and one average per acquisition. Pyruvate-to-lactate flux was quantified as the lactate/pyruvate ratio, which reflects glycolytic activity within tumors.(29)

### HMGB-1 ELISA assay

Mouse serum samples were analyzed for High Mobility Group Protein B1 (HMGB-1) levels using the Mouse HMGB-1 ELISA Kit (Thermo Fisher Scientific, Catalog No. EEL102) according to the manufacturer’s protocol. Serum samples were prepared by allowing clotting for 1 hour at room temperature, followed by centrifugation at 1000 × g for 20 minutes at 4°C to collect the supernatant. Briefly, standards and pretreated samples were added to a 8-well strip coated with HMGB-1-specific antibodies and incubated at 37°C for 90 minutes. After washing, biotinylated detection antibody and HRP-conjugate solutions were sequentially added, with incubation and washing steps in between. Substrate solution was then added, and the reaction was stopped with sulfuric acid after 15 minutes. Absorbance was measured at 450 nm using a microplate reader, and sample concentrations were determined from a standard curve.

### RNA transcriptomics

RNA extraction from 50 mg of xenograft tissue from mice treated with PD1/RT combination therapy (with or without carbogen) and subsequent transcriptomic analysis were outsourced to Novogene as a commercial service. Total RNA quality was assessed using an Agilent Bioanalyzer 2100, ensuring RNA integrity number (RIN) values above 7. Libraries were prepared using poly-A enrichment to isolate mRNA, followed by cDNA synthesis and construction of libraries with an insert size of 250–300 bp. Sequencing was performed on the Illumina NovaSeq 6000 platform, generating paired-end 150 bp reads. Quality control ensured a Q30 score exceeding 80%. Clean reads were aligned to the mouse reference genome (GRCm38/mm10) using HISAT2,(30) and gene expression levels were quantified with featureCounts v1.5.0-p3. Differential expression analysis was conducted using EdgeR v3.22.5(31) excluding noncoding RNA and pseudogenes through gene ID lookup in version 79 of the Ensembl M. musculus database. Genes with fewer than one count per million in any sample were filtered out. The final model used a generalized linear model likelihood ratio test(32) with a multiplicity-adjusted p-value threshold of 0.05 using the trimmed mean of M values (TMM) to normalize values across samples.(33) Gene Ontology (GO) analysis of differentially expressed genes was performed using PAGODA.(34)

### Statistical Analysis

The significance of differences between groups was evaluated using statistical tests selected based on data distribution. For all analyses, normality of residuals was assessed using the Anderson-Darling test (alpha=0.05). When normality assumptions were met, Student’s t-test was applied for comparisons between two groups, while differences among more than two groups were analyzed using one-way ANOVA. When data deviated from normality, Mann-Whitney U test was used for two-group comparisons and Kruskal-Wallis test for multi-group analyses. Multiple comparisons were corrected using Dunnett’s post hoc test. A two-tailed P-value <0.05 was considered statistically significant.

All statistical analyses were performed using GraphPad Prism software (version 10.4.1). Data is presented as mean ± standard error of the mean (SEM) unless otherwise specified. Sample sizes for each group were determined based on preliminary studies to ensure adequate power for detecting significant differences. Normality of data distribution was assessed prior to applying parametric tests, and non-parametric alternatives were used when data deviated from normality.

## Results

### MC38, but not B16.F10, exhibits an abscopal effect in a C57BL/6 mouse model

To create a mouse model demonstrating the abscopal effect, we established both primary and remote tumors.(16) The primary tumor, which would receive direct radiation treatment, was created by injecting 1x10^6^ MC38 tumor cells into the right hindleg. The remote tumor, representing a metastatic site that would not receive direct radiation, was established by injecting 2x10^5^ cells into the left hindleg. This approach intentionally allowed the primary tumor to grow larger, aiming to maximize tumor antigen release from the irradiated tissue, potentially triggering a systemic immune response. (16) When tumors reached approximately 100 mm³, we initiated a hypofractionated radiation treatment regimen (8 Gy/day for 3 consecutive days) to the primary tumor. This regimen was chosen to induce a significant abscopal effect, as conventional fractionated low-dose radiation can potentially induce T cell exhaustion. (35–38) Concurrently, we administered PD-1 blockade to further enhance CD8^+^ T cell activation by inhibiting its inactivation signal. Figure 1A illustrates this experimental design, while Figure 1B provides a detailed timeline of the treatment regimen.

**Figure 1.**
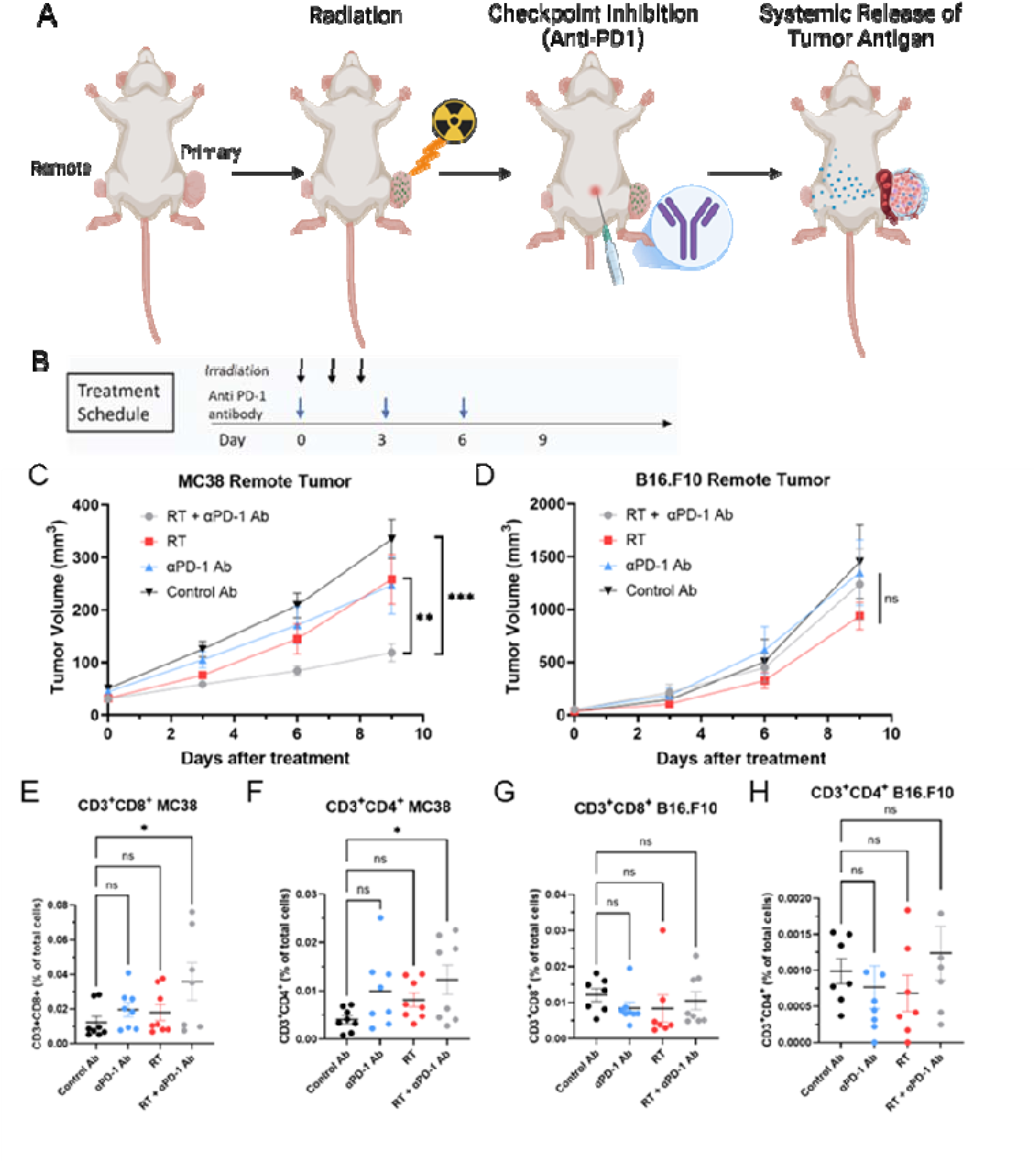
Tumor growth and immune activation following combination therapy in MC38 and B16.F10 models. (A) Schematic representation of the abscopal effect mechanism. Radiation therapy (RT) induces tumor-specific antigen release from the irradiated primary tumor, which is presented by dendritic cells to activate CD8^+^ T cells. Activated tumor-specific CD8^+^ T cells circulate systemically and target non-irradiated metastatic lesions. PD-1 blockade enhances this immune response by inhibiting the PD-1/PD-L1 pathway. (B) Timeline of the treatment schedule, including irradiation (8 Gy/day for three consecutive days) and αPD-1 antibody administration. (C-D) Tumor growth curves for remote tumors in MC38 (C) and B16.F10 (D) models following treatment with control, RT alone, PD-1 blockade alone, or combination therapy. Combination therapy significantly suppressed remote tumor growth in the MC38 model but not in the poorly immunogenic B16.F10 model. (E-H) Flow cytometry analysis of tumor-infiltrating lymphocytes (TILs) in remote tumors. In the MC38 model, combination therapy increased CD8^+^ T cell infiltration (E) and CD4^+^ T cell infiltration (F). In the B16.F10 model, no significant differences in TIL populations were observed across treatment groups (G and H). Data are presented as mean ± SEM; statistical significance is indicated (*P &<0.05, ns = not significant).

Figure 1C illustrates the progression of remote MC38 tumors following treatments with vehicle, radiation, PD-1 blockade, and the combination treatment. The primary tumor disappeared in all cases after combination treatment. Although groups subjected to either radiation therapy or PD-1 blockade monotherapy showed a relatively slower tumor growth compared to the control group, no statistically significant differences were observed among these groups at day 9 after treatment. This outcome is likely due to the systemic immune activation triggered by radiation therapy-induced tumor antigens or PD-1 blockade. On the other hand, the sole group receiving the combination treatment of PD-1 blockade and radiation therapy showed clear suppression of remote tumor growth with reduced variability within the group, suggesting successful induction of the abscopal effect. The same set of experiments were conducted on B16.F10 tumors (Figure. 1D). However, these tumors did not exhibit the abscopal effect observed in the MC38 tumor model.

### Flow Cytometry Confirms Enhanced T Cell Infiltration in Remote MC38 Tumors Following **α**-PD-1/Radiation Combination Therapy

To verify the relationship between the abscopal effect and immune activation within remote tumors, flow cytometric analysis was performed using cell suspensions obtained from both MC38 and B16.F10 tumor samples in each treatment group. Notably, in the combination group of MC38 tumors, the proportion of CD8^+^ cells within the total cell population was the highest compared to the other groups, indicating an increased overall number of tumor-infiltrating CD8^+^ T cells (Figure 1E). A similar trend was observed in CD4^+^ T cells (Figure 1F). In contrast, such robust T cell activation was not observed in B16.F10 tumors (Figure 1G and 1H). Within the combination group, CD8^+^ T cell activation is more pronounced within MC38 tumors, aligning with the observed differences in the abscopal effect. This suggests the lack of an abscopal effect in B16.F10 may be related to the differences in PD-1 immunogenicity between MC38 and B16.F10, (37,38) but other factors such as the significantly faster growth rate of B16.F10 may also be involved.

### Increases in Hyopoxic Fraction and Extracellular Volume, Enhanced Glycolysis in Tumors Exhibiting the Abscopal Effect

We used MRI-based multimodal imaging techniques to assess metabolic and functional changes in remote tumors across all treatment groups. On the 9th day post-treatment, we performed EPR oximetry to generate 3D in vivo pO2 maps of the tumors. Figure 2A displays representative pO2 maps for each treatment group. While median pO2 levels were similar across groups, the hypoxic fraction below 10 mmHg (HF10) was significantly lower in the combination treatment group (Figure 2B). HF10, often used to assess tumor hypoxia or radiosensitivity, suggests either enhanced tumor perfusion or reduced oxygen consumption in the combination treatment group. To further investigate these changes, we conducted dynamic contrast-enhanced (DCE) MRI using intravenous Gd-DTPA injection. Figure 2C shows representative images of temporal changes in Gd-DTPA concentration for each group, while Figure 2D illustrates the time-intensity kinetics of Gd-DTPA. The time to peak remained consistent across groups, indicating similar perfusion characteristics. However, the Area Under the Curve at 10 minutes (AUC10min) increased in the combination treatment group (Figure 2F). AUC10min reflects the extracellular volume at equilibrium, potentially indicating increased cell death or reduced tumor cellularity in response to the combination treatment.

**Figure 2.**
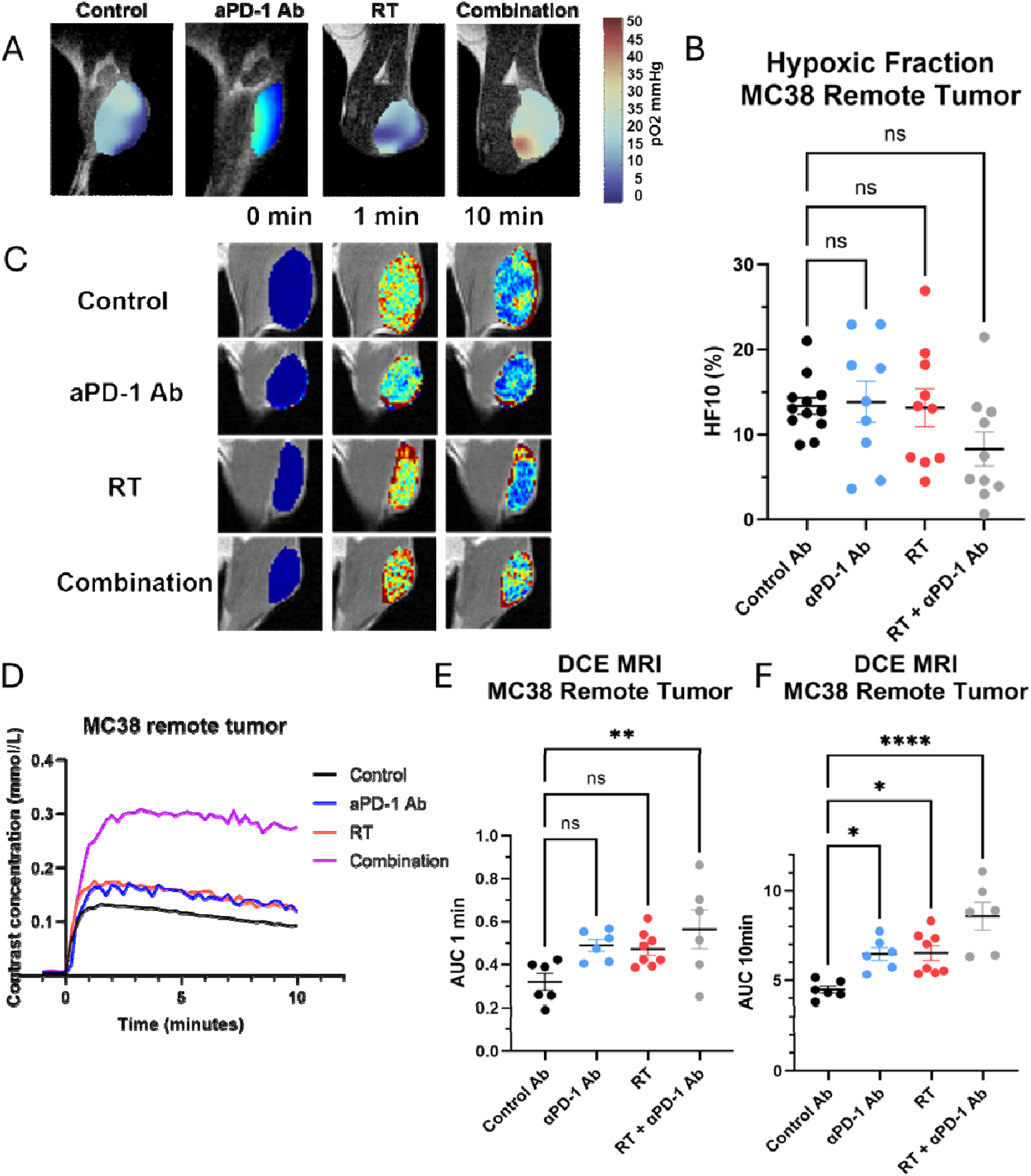
Tumor oxygenation and perfusion characteristics in the remote tumor following combination therapy with PD-1 blockade and radiation. (A) Representative pO^2^ maps of MC38 tumors from each treatment group (control, anti-PD-1 antibody, radiation therapy [RT], and combination therapy) generated using EPR oximetry. (B) Quantification of the hypoxic fraction (HF10), defined as the percentage of tumor volume with pO₂ &<10 mmHg. (C) Representative DCE-MRI images showing temporal changes in Gd-DTPA contrast concentration (0 min, 1 min, and 10 min post-injection) for each treatment group. (D) Time-intensity curves of Gd-DTPA uptake in MC38 tumors. (E-F) Quantification of perfusion metrics from DCE-MRI: AUC1min (E) and AUC10min (F). Both metrics were significantly elevated in the combination therapy group, reflecting enhanced perfusion and permeability. Data are presented as mean ± SEM; statistical significance is indicated (*P &<0.05, **P &<0.01, ***P &<0.001).

To confirm these findings, we conducted diffusion-weighted MRI and hyperpolarized 13C-pyruvate MRI studies across all four treatment groups to evaluate changes in tumor cellularity and glycolysis associated with the abscopal effect. Diffusion-weighted MRI was used to calculate the Apparent Diffusion Coefficient (ADC), which reflects the microscopic motion of water molecules. Lower ADC values typically indicate higher cellularity due to restricted water diffusion. In the combination treatment group, where the abscopal effect was observed, ADC values were significantly higher compared to the control group (Figure 3A, 3B). This suggests decreased cellularity in the combination treatment group, likely due to the cytotoxic abscopal effect, contrasting with increased cellularity in the control tumors due to continued progression. Hyperpolarized 13C-pyruvate MRI was employed to monitor pyruvate-to-lactate flux, indicating the extent of glycolytic activity in the tissue. This technique uses exogenously administered hyperpolarized 13C-pyruvate to generate spectrally resolved metabolite peaks. Representative images are shown in Figure 3C. The lactate/pyruvate ratio, an indicator of glycolytic flux, exhibited a modest, statistically non-significant decrease in the combination treatment group compared to the control group (Figure 3D). This slight reduction in glycolytic activity could be associated with improved perfusion or a minor decrease in oxygen consumption linked to the abscopal effect, potentially reducing competition for glucose and oxygen between tumor cells and immune cells. The changes were subtle, likely reflecting the limited efficacy of the abscopal effect and the inherently glycolytic nature of activated immune cells. These findings align with previous research on metabolic changes following immune checkpoint blockade.(24)

**Figure 3.**
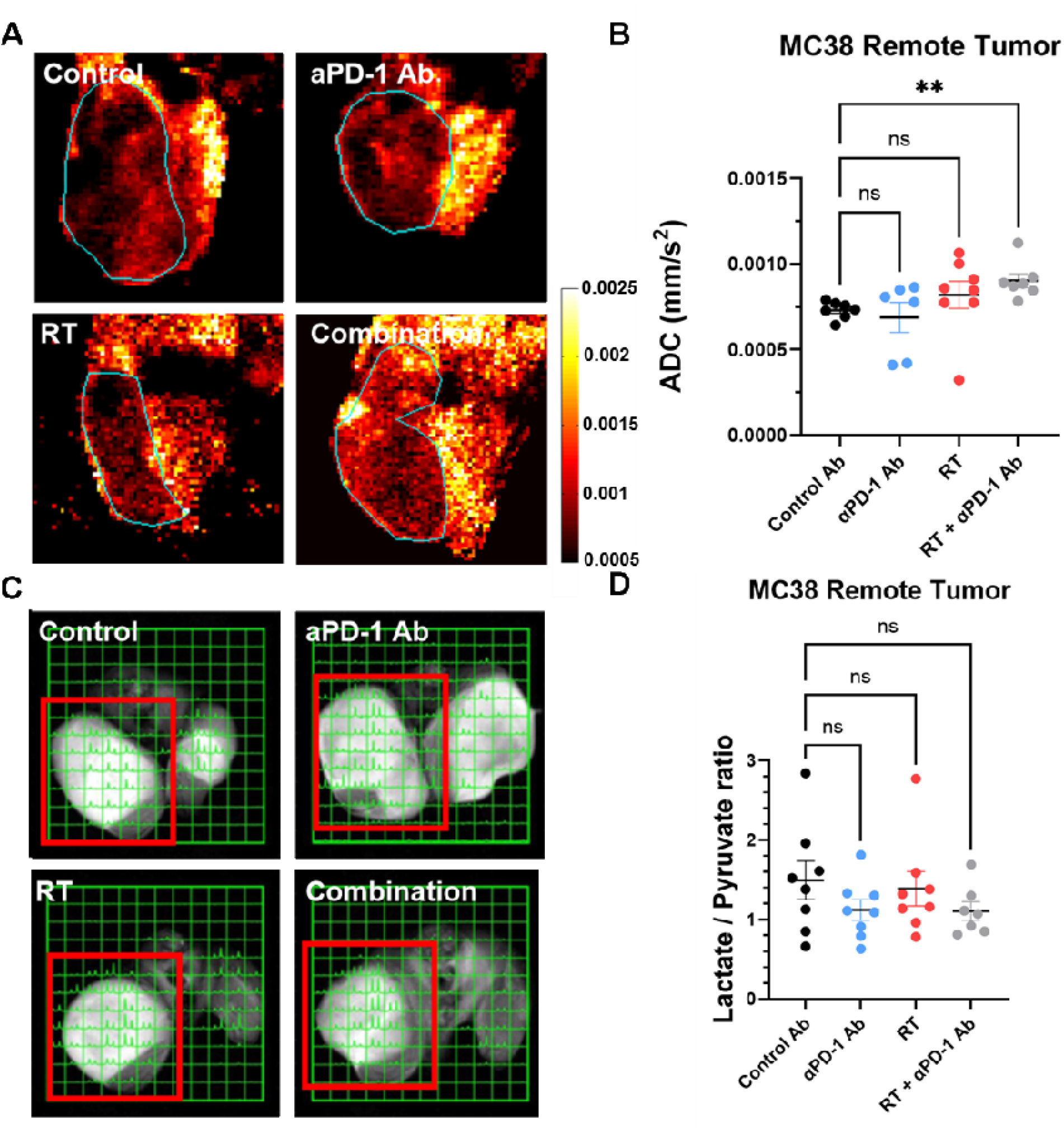
**Treatment-induced changes in tumor cellularity and glycolytic activity.**(A) Representative apparent diffusion coefficient (ADC) maps of MC38 tumors from each treatment group (control, anti-PD-1 antibody, radiation therapy [RT], and combination therapy). (B) Quantification of ADC values in MC38 remote tumors. Combination therapy significantly increased ADC values compared to the control group, indicating reduced cellularity due to treatment-induced cell death (**P &<0.01). (C) Representative hyperpolarized ^13^C MRI images showing lactate and pyruvate signals in MC38 tumors for each treatment group. (D) Quantification of the lactate/pyruvate ratio in MC38 remote tumors. Data are presented as mean ± SEM, with statistical significance indicated (*P &<0.05, **P &<0.01).

### Hypoxia and Reduced Extracellular Volume in the Primary Tumor Predict Future Growth in the Remote Tumor

Tumors exhibiting the abscopal effect in response to combination therapy showed increased extracellular volume, decreased cellularity, and reduced glycolytic activity, suggesting these imaging biomarkers could potentially serve as indicators for assessing the abscopal effect in metastatic tumors treated with combination immunotherapy and radiation. To identify predictive biomarkers for the abscopal effect, we conducted EPR oximetry and DCE MRI prior to combination therapy, recording biomarkers from both primary and metastatic tumors (Figure 4A). Notably, these biomarkers in remote tumors did not correlate with the size of the remote tumor on day 9, indicating limited relevance to anti-tumor immune activity following combination therapy (Supplementary Figure 1A and 1B). However, we observed a robust negative correlation between perfusion and permeability markers in the primary tumor and the size of the remote tumor on day 9 (Figure 4B and 4C). Additionally, the hypoxic fraction (HF10, pO_2_<10 mm Hg) showed a positive correlation with metastatic tumor size, albeit to a lesser extent (Figure 4D). These findings suggest that high tumor perfusion and permeability in the primary tumor before combination treatment may predict a stronger induction of the abscopal effect. Importantly, radiation sensitivity is unlikely to contribute to these results, as the primary tumor disappeared in all cases after combination treatment.

**Figure 4.**
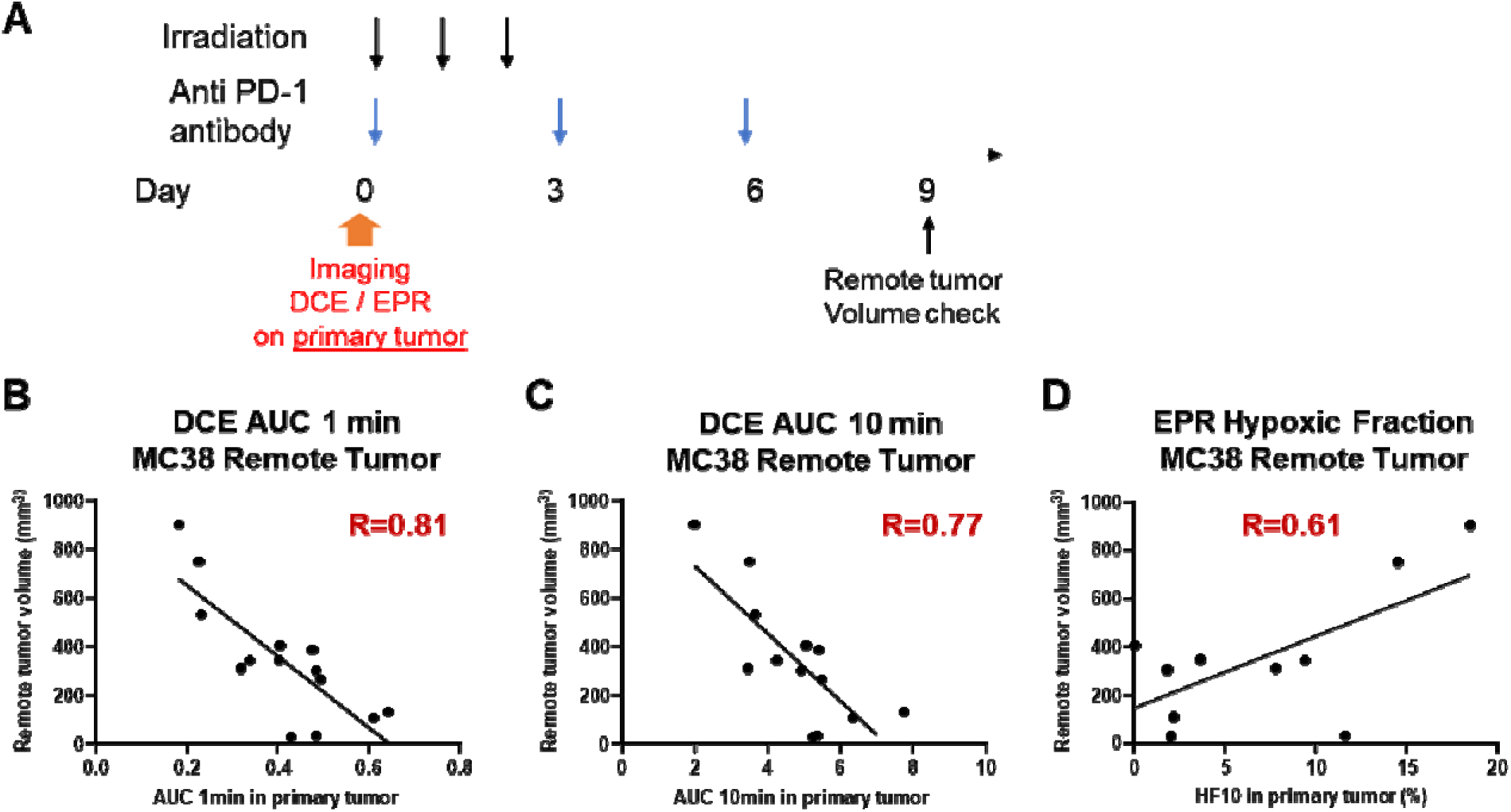
Tumor perfusion and oxygenation in the primary tumor predicts future growth in the remote tumor in the MC38 model. (A) Timeline of treatment and imaging schedule. DCE-MRI and EPR oximetry were performed on the primary tumor prior to combination therapy (anti-PD-1 antibody and irradiation). Remote tumor volume was measured on day 9 post-treatment to assess systemic responses. (B-D) Correlations between primary tumor imaging metrics on the primary tumor on day 1 and remote tumor volume on day 9. AUC1min (B) and AUC10min (C) from DCE-MRI inversely correlate with future remote tumor volume, indicating that higher early perfusion and increased extracellular volume in the primary tumor predict better systemic control of remote tumors (D) Hypoxic fraction (HF10) from EPR oximetry positively correlates with remote tumor volume, demonstrating that higher hypoxia in the primary tumor predicts poorer systemic outcomes.

### Carbogen Enhances the Abscopal Effect

To investigate the role of improved perfusion in achieving a successful abscopal effect, we compared tumor growth between two groups: one receiving combination therapy and another receiving combination therapy with enhanced perfusion preconditioning. In the latter group, tumor perfusion was significantly increased by exposing mice to carbogen (5% carbon dioxide + 95% oxygen) during irradiation.(39) The vasodilatory effect of 5% carbon dioxide counteracted the vasoconstrictive effect of high oxygen concentration, mimicking primary tumors with high perfusion and oxygenation that exhibited a stronger abscopal effect in remote tumors (Figure 4B-D). (40,41) To minimize potential long-term effects, carbogen exposure was limited to 120 minutes during irradiation (40 minutes x 3 radiation therapies). DCE MRI confirmed the enhancing effect of carbogen on perfusion, showing a 71% increase in AUC1min and a 49% increase in AUC10min (Figure 5A and 5B). The results of the tumor growth comparison are shown in Figure 5C. Mice exposed to carbogen during combination therapy showed suppressed growth of remote tumors compared to those exposed to air in the MC38 tumor model, indicating an augmented abscopal effect. Notably, this enhanced effect was observed even when tumors reached approximately 100 mm³, a stage where combination treatment typically becomes less effective due to rapid tumor growth. In contrast, the B16.F10 model showed no discernible difference in tumor growth between groups, suggesting that carbogen alone does not suppress tumor growth when combination treatment fails to induce sufficient immune activation (Figure 5D).

**Figure 5.**
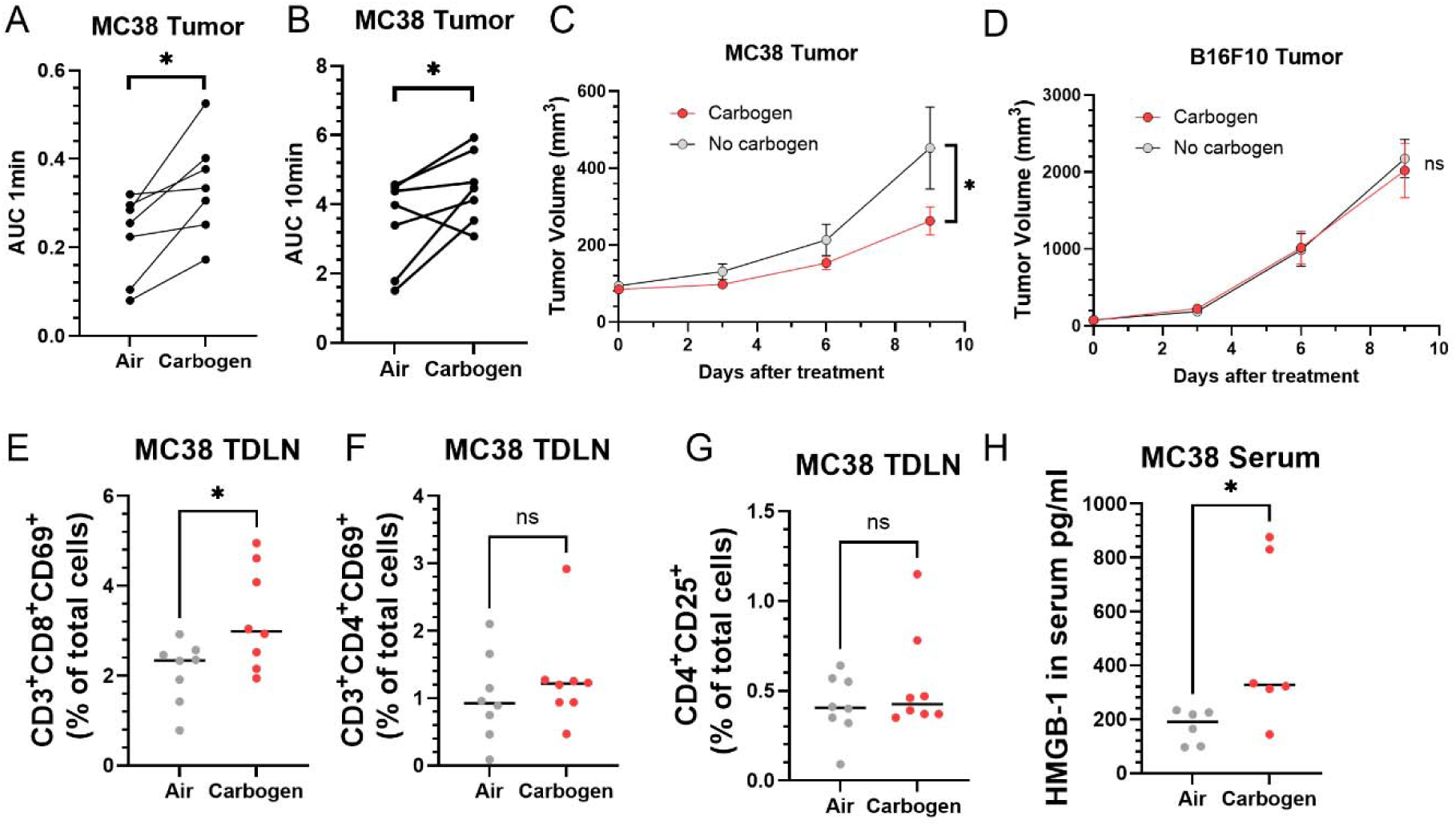
**Carbogen exposure enhances tumor perfusion, immune activation, and systemic anti-tumor responses in the MC38 model**. (A-B) DCE-MRI analysis of primary tumors shows that carbogen exposure significantly increases perfusion metrics, with a 71% increase in AUC1min (A) and a 49% increase in AUC10min (B) compared to air-exposed controls. (C-D) Tumor growth curves for MC38 (C) and B16.F10 (D) models following combination therapy with or without carbogen exposure. In the MC38 model, carbogen exposure significantly suppresses remote tumor growth compared to air exposure, while no significant differences are observed in the poorly immunogenic B16.F10 model. (E-G) Flow cytometry analysis of tumor-draining lymph nodes (TDLNs) in the MC38 model shows a significant increase in activated CD8+ T cells (CD3+CD8+CD69+) in carbogen-treated mice (E), with no significant changes observed in activated CD4+ T cells (F) or regulatory T cells (G).(H) Serum levels of HMGB-1, a damage-associated molecular pattern (DAMP), are significantly elevated in carbogen-treated mice, suggesting enhanced immune activation.

**Figure 6.**
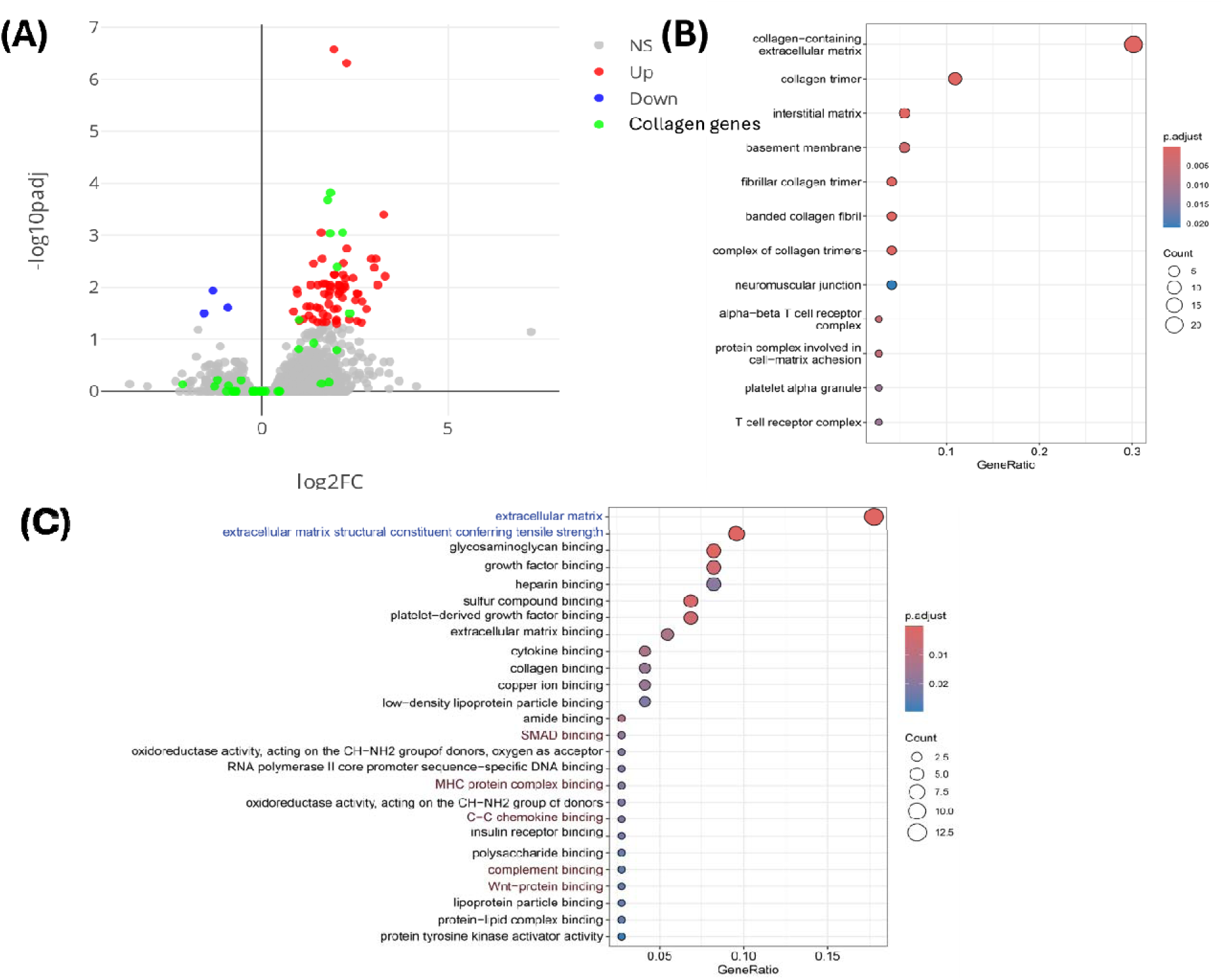
Transcriptomic analysis of the effect of carbogen on anti-PD-1/RT treatment. Volcano plot showing differentially expressed genes (DEGs) in primary tumors treated with combination therapy (anti-PD-1 antibody and radiation) with or without carbogen exposure. DEGs were defined as genes with an adjusted P-value &<0.05 and |log2 fold change| > 1. Red dots represent genes upregulated in response to carbogen treatment, blue dots represent downregulated genes, and green dots highlight collagen-related genes. (B) Gene Ontology (GO) enrichment analysis of upregulated DEGs. (C) GO molecular function analysis of upregulated DEGs.

To confirm whether carbogen exposure during radiation treatment enhances immune activation, we examined the primary tumor-draining lymph node using flow cytometry 4 days after treatment. In MC38 tumors, we observed a significant increase in the fraction of activated CD8^+^ T cells (CD69 positive) in the total live cell population in the carbogen-exposed group compared to the air-exposed group (Figure 5E). However, no differences were found in activated CD4^+^ T cells and regulatory T cells between the two groups (Figures 5F and 5G). This suggests that carbogen exposure augments the activation of cytotoxic T cells following combination treatment.

The enhanced abscopal effect observed with carbogen exposure is attributed to the systemic circulation of activated cytotoxic T cells, which target remote tumors. We propose that carbogen exposure enhances immune activation by facilitating the effective release and uptake of tumor antigens and immune-associated cytokines. Specifically, lethal radiation damage triggers the release of HMGB-1 from tumor cells, leading to complex immune responses.(42–44) Previous studies have shown that the activation of anti-tumor immune responses occurs when HMGB-1 release surpasses a specific threshold. In our study, we found a 2.5-fold increase in serum HMGB-1 levels in mice treated with combination therapy and carbogen exposure compared to those without carbogen exposure (Figure 5H). This provides a partial explanation for the augmented abscopal effect observed with carbogen exposure.

To further investigate the molecular mechanisms underlying the enhanced abscopal effect observed with carbogen exposure, we performed RNA sequencing (RNA-seq) experiments on tumor samples from the αPD-1/radiation therapy treatment groups with and without carbogen breathing. The analysis was conducted 9 days post-treatment to capture longer-term transcriptional changes. After filtering out non-coding and lowly expressed genes, we identified 68 upregulated and 3 downregulated genes in the carbogen-exposed samples compared to the control group. Interestingly, many of the genes typically associated with the abscopal effect,(5) such as interferons and cytokines, were not significantly differentially expressed, likely due to the time elapsed since treatment. However, we observed a notable upregulation of collagen-related genes. Gene Ontology (GO) function analysis revealed a strong association with collagen-related extracellular remodeling, with T cell receptor and complement system activation to a lesser extent. These findings suggest a prominent fibroblast invasion in the aftermath of treatment, potentially indicating ongoing tissue remodeling and immune response modulation in the tumor microenvironment

## Discussion

The abscopal effect, where localized radiation therapy induces regression of distant metastatic lesions, represents a promising approach for treating metastatic cancer through systemic immune activation. This phenomenon fundamentally depends on the ability of CD8^+^ T cells to recognize and attack distant tumors following local radiation therapy, as demonstrated by increased CD8^+^ T cell infiltration and activation in tumor-draining lymph nodes of responding tumors. However, its clinical utility has been limited by inconsistent occurrence, even when radiation is combined with immune checkpoint inhibitors. While various factors influence this variability, the tumor microenvironment (TME) determines treatment outcomes through its effects on immune cell trafficking, function, and distribution.

Among these factors affecting immune responses, tumor perfusion fundamentally shapes immune cell trafficking and function within the tumor microenvironment. Enhanced perfusion facilitates the distribution of immune cells, cytokines, and other immune mediators throughout the tumor tissue, while also supporting the metabolic requirements of activated immune cells. Recent studies have demonstrated the importance of perfusion in immunotherapy response. In melanoma brain metastases, lower plasma volume (rVp) on DCE-MRI distinguished pseudoprogression from true progression, indicating that vascular characteristics influence immune responses.(45) Similarly, in metastatic melanoma patients receiving checkpoint inhibitors, successful treatment responses were associated with higher baseline vascular transfer constant (Ktrans) and extracellular volume (ve), suggesting that better perfusion supports immune cell function.(46) This relationship between perfusion and immune function suggests that perfusion metrics could serve as surrogate markers for a tumor’s capacity to mount effective immune responses.

Given the central role of perfusion in immune responses, we hypothesized that vascular characteristics might shape abscopal responses. We established a dual-tumor model in mice,(16) with a larger primary tumor in the right hind leg to enhance antigen presentation during radiotherapy (8 Gy × 3 consecutive days) and a smaller, unirradiated remote tumor in the left hind leg to assess systemic responses. Anti-PD-1 antibody was administered on days 0, 3, and 6. This design leverages two key mechanisms: radiation-induced immunogenic cell death releasing tumor antigens for dendritic cell capture, and PD-1 blockade preventing T cell exhaustion. The approach proved successful in the immunogenic MC38 model, where combination therapy significantly suppressed remote tumor growth and increased CD8^+^ T cell infiltration. In contrast, the less immunogenic B16.F10 model showed no response.

In the immune-sensitive MC38 model, abscopal responses also led to notable changes in the *remote* tumor microenvironment. EPR oximetry showed reduced hypoxia in remote tumors of mice receiving combination therapy, with a marked decrease in the hypoxic fraction (HF10) compared to control groups. Using DCE-MRI, we observed enhanced perfusion in remote tumors after combination therapy, as shown by greater contrast agent uptake in both early (AUC1min) and late (AUC10min) phases. These vascular improvements were coupled with decreased tumor cellularity, as indicated by elevated ADC values. The time-intensity curves revealed enhanced contrast agent concentrations in the combination therapy group compared to single treatments or controls, suggesting better vascular function.

None of these changes in the tumor microenvironment in the remote tumor showed prognostic value for treatment response. However, surprisingly, vascular characteristics of the *primary* tumor before treatment showed significant prognostic value for treatment response in metastatic lesion models, despite the primary tumor disappearing after treatment in every case. Higher perfusion (AUC1min) and extracellular volume (AUC10min) in primary tumors correlated with better control of remote lesions, while increased hypoxia in the primary tumor predicted poor systemic responses. While combination therapy induced clear changes in the remote tumor microenvironment - including increased extracellular volume (AUC10min), decreased cellularity (ADC), and reduced hypoxia - these changes did not predict treatment outcomes. Instead, the predictive power resided exclusively in the primary tumor’s pre-treatment characteristics. This distinction is crucial: despite observable changes in remote tumors, only primary tumor vascular metrics before treatment reliably predicted systemic responses. This finding suggests that initial immune priming in the primary tumor, rather than pre-existing conditions in remote tumors, drives systemic responses to combination therapy.

The correlation between primary tumor vascular characteristics and systemic treatment response points to a mechanistic cascade linking vascular function to systemic immune responses. While improved oxygenation can enhance radiation sensitivity, our data suggest that perfusion-mediated immune activation drives the abscopal effect. In the MC38 model, enhanced perfusion during radiation therapy increased the release and distribution of tumor antigens, as shown by elevated serum HMGB-1 levels. This led to increased antigen-presenting cell maturation and subsequent CD8^+^ T cell activation in tumor-draining lymph nodes. Notably, primary tumors regressed completely with the 8 Gy × 3 radiation protocol regardless of perfusion status, suggesting that enhanced immune activation, rather than direct radiation effects, may mediate improved systemic responses.

Assessing primary tumor vasculature before treatment could guide therapy selection, with poorly perfused tumors potentially benefiting from strategies to enhance vascular function before starting combination therapy. To test this hypothesis, we exposed mice to carbogen (95% O2 + 5% CO2) during radiation treatment. Carbogen exposure significantly enhanced tumor perfusion, as measured by DCE-MRI, with a 71% increase in AUC1min and a 49% increase in AUC10min compared to air-exposed controls. This enhanced perfusion translated to improved systemic responses - MC38 tumor-bearing mice exposed to carbogen during combination therapy showed significantly suppressed growth of remote tumors compared to air-exposed controls. Importantly, this effect was specific to the immunogenic MC38 model, as the poorly immunogenic B16.F10 model showed no response to carbogen exposure.

The enhanced systemic response correlated with increased immune activation in tumor-draining lymph nodes, specifically a higher fraction of activated CD8^+^ T cells (CD69^+^) in carbogen-treated mice. This selective activation of cytotoxic T cells, without changes in CD4^+^ T cells or regulatory T cells, suggests that improved perfusion specifically enhances cytotoxic immune responses. Supporting this mechanism, serum levels of HMGB-1, a key damage-associated molecular pattern, were 2.5-fold higher in carbogen-treated mice. While we used carbogen to enhance perfusion in this proof-of-concept study on the basis of the successful ARCON trials,(47,48) other clinically available vasodilators could potentially achieve similar effects, offering multiple approaches for clinical translation.

Several limitations should be acknowledged in this study. The use of two representative cell line models, the immunogenic MC38 cell line, and the less immunogenic B16.F10 line might limit generalizability. Further studies with diverse models are necessary to confirm the reproducibility of the results. Additionally, to enhance blood perfusion in cancer, temporary carbogen exposure was employed and confirmed by DCE MRI. However, modulating perfusion can be complicated by the reaction of blood vessels to vasodilation, including the “blood stealing phenomenon” resulting from differences in vessel constriction or dilation ability.(49) Tumor vasculature, often immature, may lack the ability to react to vasodilators, leading to paradoxical hypoperfusion. Therefore, when implementing the proposed enhanced perfusion strategy in preclinical or clinical settings, it will be crucial to monitor its effect on tumor vessels in advance.

## Conclusion

In conclusion, our findings establish perfusion as a critical determinant of the abscopal effect in two key ways. First, primary tumor perfusion metrics before treatment serve as predictive biomarkers for systemic responses, offering a potential tool for patient stratification. Second, enhancing perfusion through carbogen exposure during radiation therapy augments immune activation and improves treatment outcomes in immunogenic tumors. While EPR oximetry remains confined to preclinical studies, DCE-MRI metrics provide immediately available clinical tools for assessing tumor perfusion. The combination of perfusion-based patient selection and vascular modification strategies offers a promising approach to enhance the consistency and efficacy of combination immunotherapy and radiation treatment in metastatic cancer. Future studies should focus on validating these findings across diverse tumor types and exploring alternative methods to enhance tumor perfusion while carefully monitoring vascular responses.

## Acknowledgements

We thank Daniel R. Crooks and Cristopher Ricketts for their helpful discussion.

## Author Contributions

Kota Yamashita: Investigation, Methodology, Writing – original draft, Yu Saida: Investigation, Methodology. Yasunori Otowa: Investigation, Takeshi Ito: Investigation, Kazutoshi Yamamoto: Investigation. Hellmut Merkle: Resources, Gadisetti VR Chandramouli: Formal analysis, Investigation, Software. Nallathamby Devasahayam: Resources, Investigation. James B. Mitchell: Project administration, Supervision, Funding acquisition. Murali C. Krishna: Conceptualization, Project administration, Methodology, Supervision, Funding acquisition. Jeffery R. Brender: Investigation, Methodology, Formal analysis, Writing – original draft, Writing – review & editing. Shun Kishimoto: Investigation, Conceptualization, Methodology, Supervision, Writing – original draft, Writing – review & editing.

