## Supplementary Figure 1 for "Improved Tumor Blood Flow Enhances the Abscopal Effect: Preclinical Assessment in Mice Treated with Combined Radiation and PD-1 Blockade Therapy"

A

AUC10min Remote day0 vs Remote Tumor Volume day9

| Permeability in remote tumor at day0 | remote tumor volume |
| --- | --- |
| 6.30314 | 110.466 |
| 5.86747 | 32.49 |
| 5.77335 | 36.1415 |
| 4.77053 | 417.15 |
| 5.82555 | 408.726 |
| 5.60266 | 305.809 |
| 4.10597 | 348.8 |
| 5.11477 | 752.328 |
| 3.7255 | 313.326 |
| 2.02983 | 903.96 |
| 3.33243 | 346.8 |

B

HF10 Remote day0 vs Remote Tumor Volume day 9

| HF10 in remote tumor at day0 | remote tumor volume |
| --- | --- |
| 7.8285 | 110.466 |
| 15.1631 | 32.49 |
| 1.8398 | 36.1415 |
| 6.3291 | 417.15 |
| 0 | 408.726 |
| 1.7137 | 305.809 |
| 2.8074 | 348.8 |
| 23.4247 | 752.328 |
| 15.1877 | 313.326 |
| 15.8519 | 903.96 |
| 19.6198 | 346.8 |

Supplementary Figure 1: Correlations between (A) DCE AUC 10min and (B) EPR Hypoxic Fraction on the remote tumor on day 0 and remote tumor volume on day 9.
